# Homeostats – the hidden rulers of ion homeostasis in plants

**DOI:** 10.1101/2024.04.15.589567

**Authors:** Ingo Dreyer, Naomí Hernández-Rojas, Yasnaya Bolua-Hernández, Valentina de los Angeles Tapia-Castillo, Sadith Zobeida Astola-Mariscal, Erbio Díaz-Pico, Franko Mérida-Quesada, Fernando Vergara-Valladares, Oscar Arrey-Salas, María Eugenia Rubio-Meléndez, Janin Riedelsberger, Erwan Michard

**Affiliations:** Electrical Signaling in Plants (ESP) Laboratory–Centro de Bioinformática y Simulación Molecular (CBSM), Facultad de Ingeniería, Universidad de Talca, 2 Norte 685, Talca CL-3460000, Chile; Programa de Doctorado en Ciencias mención Biología Vegetal y Biotecnología, Universidad de Talca, 2 Norte 685, Talca CL-3460000, Chile; Programa de Magíster en Bioquímica y Biología Molecular, Universidad de Talca, 2 Norte 685, Talca CL-3460000, Chile; Programa de Magíster en Hortofruticultura, Universidad de Talca, 2 Norte 685, Talca CL-3460000, Chile; Programa de Doctorado en Ciencias mención Modelado de Sistemas Químicos y Biológicos, Universidad de Talca, 2 Norte 685, Talca CL-3460000, Chile; Instituto de Ciencias Biológicas, Universidad de Talca, Campus Talca, Avenida Lircay, Talca CL-3460000, Chile

**Keywords:** homeostasis, transporter networks, modeling, membrane transport, thermodynamics, quantitative biology

## Abstract

Ion homeostasis is a crucial process in plants that is closely linked to the efficiency of nutrient uptake, stress tolerance and overall plant growth and development. Nevertheless, our understanding of the fundamental processes of ion homeostasis is still incomplete and highly fragmented. Especially at the mechanistic level, we are still in the process of dissecting physiological systems to analyze the different parts in isolation. However, modeling approaches have shown that it is not individual transporters but rather transporter networks (homeostats) that control membrane transport and associated homeostatic processes in plant cells. To facilitate access to such theoretical approaches, the modeling of the potassium homeostat is explained here in detail to serve as a blueprint for other homeostats. Based on a few, elementary knowledge about the thermodynamics of the different transport processes, it is possible to draw fundamental conclusions about the properties and physiology of the transporter network.

## Introduction

Homeostasis is the ability of an organism to maintain stable physico-chemical conditions that are compatible with cell metabolism despite fluctuations in the external environment. Plants are subject to permanent changes in the availability of water, nutrients and light energy as well as other parameters, such as temperature, which influence the entire metabolism. Plants are a semi-open system in which the concentrations of apoplastic fluid vary and depend on environmental parameters, in particular the water potential of the soil and the concentration of dissolved substances, on the one hand, and on internal factors, such as the opening of the stomata, on the other. Ion homeostasis is central to plant physiology as it regulates and maintains cell turgor, energizes and controls nutrient flux (especially of sugars and amino acids) and is at the heart of numerous signaling pathways dependent on Ca^2+^ and H^+^. Each individual plant cell contributes to ion homeostasis by maintaining the concentrations of free ions in the syncytium and tissues that are optimal for growth and efficient ion-dependent signaling. Given the importance of ion homeostasis, understanding the mechanisms involved is crucial for improving crop yields and for developing strategies to cope with environmental stresses such as drought and nutrient deficiency.

Ion homeostasis involves the transport and sequestration or release into or out of various compartments such as apoplast, cytosol, vacuole, mitochondria, plastids, endocytic and exocytic membrane complexes. All these membranes have their own specific transporters and associated regulatory factors (regulatory proteins, voltage, pH, pCa, etc.) that form a highly complex system. Before approaching such complexity, a first step towards understanding the principles of ion homeostasis in plants can be a theoretical approach, looking at the simplest possible system and searching for emergent properties. Over the last four to five decades, our knowledge of plant membrane transport has improved considerably thanks to the development of new techniques and molecular genomic analyses (Ward et al., 2009; Hedrich, 2012; Anschütz et al., 2014; Jegla et al., 2019; Stanton et al., 2022; Blatt, 2024) and physiological data that enable such a theoretical approach. Nevertheless, membrane transport in plants is still widely analyzed using more than 70 years old concepts. For instance, the classification of transporters as “high-affinity” and “low-affinity” transporters was very helpful in the 1950s and 1960s in identifying different transporters *in vivo* and to distinguishing between them (Epstein and Hagen, 1952; Epstein et al., 1963). However, the categorization into these two groups and the interpretation that transporters with “high affinity” are active at low concentrations, while transporters with “low affinity” take over at higher concentrations, led to unclear conclusions and even contradictions (for further details, please see Dreyer, 2017; Dreyer and Michard, 2020). It became evident that there is an urgent need for a new solid theoretical foundation that not only describes the observed phenomena, but can also clearly explain them based on first principles. One such approach resulted in the theory of homeostats (Dreyer, 2021a), i.e. transporter networks that act together and exhibit dynamic properties as a system that go beyond of those of the isolated transporters (Dreyer et al., 2022; Li et al., 2024). This unbiased, systemic approach combines thermodynamics with biophysics and translates biological phenomena into the language of mathematics, enabling the derivation of analytical solutions or computational simulations of specific situations.

To increase the understanding and accessibility of such an approach, this article intends to serve as a hands-on tutorial for modeling a homeostat using the example of the potassium (K) homeostat. The transporter network of the K homeostat is composed of K^+^ channels, H^+^/K^+^ symporters, H^+^/K^+^ antiporters and H^+^-ATPases establishing the necessary proton and voltage gradients that energize the different transport processes (**Figure 1**). The system, in which the K homeostat is embedded, is a membrane that separates two compartments (internal/inside and external/outside) and is characterized by the following parameters (**Table 1**): volumes of the internal and external compartment (*Vol*_*in*_, *Vol*_*out*_), proton and potassium concentrations in the internal and external compartment ([*H*^*+*^]_*in*_, [*H*^*+*^]_*out*_, [*K*^*+*^]_*in*_, [*K*^*+*^]_*out*_), the membrane capacitance and membrane voltage (*C, V*), as well as the activities of the transporters.

**Table 1.** Parameters of the K homeostat system. The following list of parameters determines the system of a K homeostat embedded in a membrane that separates two compartments. Appropriate definitions for the parameters marked with an asterisk (*) will be provided in the course of this study.

|  |  |
| --- | --- |
| Internal compartment | $Vol_{in}$ (pL): internal Volume<br>$[K^+]_{in}$ (μM): internal $K^+$ concentration |
|  | • $[H^+]_{in}$ ($\mu M$): internal proton concentration |
| Membrane | • $C$ (pF): Membrane capacitance<br>• $V$ (mV): membrane voltage<br>• activity of $H^+$-ATPases *<br>• activity of $K^+$ channels *<br>• activity of $H^+/K^+$ symporters *<br>• activity of $H^+/K^+$ antiporters * |
| External compartment | • $Vol_{out}$ (pL): external Volume<br>• $[K^+]_{out}$ ($\mu M$): external $K^+$ concentration<br>• $[H^+]_{out}$ ($\mu M$): external proton concentration |
| Derived parameters | • $E_K = \frac{RT}{F} \cdot \ln \left( \frac{[K^+]_{out}}{[K^+]_{in}} \right)$ (Nernst potential for $K^+$; mV)<br>• $E_H = \frac{RT}{F} \cdot \ln \left( \frac{[H^+]_{out}}{[H^+]_{in}} \right)$ (Nernst potential for $H^+$; mV) |

**Figure 1.**
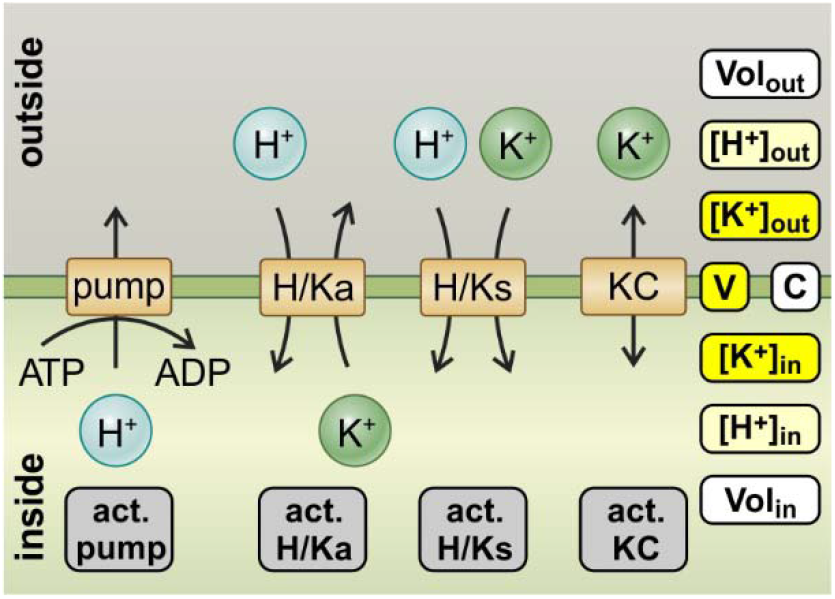
The K homeostat. The K homeostat is network of transporters that transport K^+^, in combination with the energizing proton ATPase. It consists of H^+^ ATPases (pump), H^+^/K^+^ antiporters (H/Ka), H^+^/K^+^ symporters (H/Ks) and K^+^ channels (KC), which are embedded in a membrane that separates an internal compartment (*inside*) from an external compartment (*outside*). The system is determined by 12 parameters: Three of them (*white*) are determined by the geometry of the system: internal volume (*Vol*_*in*_), external volume (*Vol*_*out*_) and the membrane capacitance (*C*). Four of them (*grey*) describe the activity of the transporters: activity of the pump (*act. pump*), the antiporter (*act. H/Ka*), the symporter (*act. H/Ks*) and the K^+^ channel (*act. KC*). Two of them ([H^+^]_in_ and [H^+^]_out_, *light yellow*) are partially influenced by buffer reactions and transport processes different from the K homeostat. The remaining three ([K^+^]_in_, [K^+^]_out_, and *V, yellow*) depend on the setting of the K homeostat and are controlled by it.

By having chosen the K homeostat, we follow a historical pattern, because research on K^+^ transport in plants has been a constant driving force in the past for progress in membrane transport and has influenced research on nutrient transport in general (Britto et al., 2021; Dreyer, 2021b). We start with explaining the mathematical description of transmembrane transport processes through the different transporters. Then we present the differential equations that describe the changes in concentrations and the membrane voltage due to the transport processes. Thereafter, we analyze the system in steady state. In a follow-up study, we will consider deviations from the steady state condition in order to get an idea about the dynamic properties of the homeostat. The different steps can in principle be applied to any other homeostat, as recently demonstrated for the auxin homeostat in plants (Geisler and Dreyer, 2024).

## Methods

### Mathematical Description of Transport Processes

A theoretical/mathematical description of membrane transport requires to quantify the transport processes involved. For this purpose, the biophysical knowledge on transport (Hille, 2001) needs to be translated into the universal language of mathematics. Here, only basic knowledge on the membrane transport is often sufficient to obtain rather simple but very powerful equations. The art of mathematical modeling lies in the elimination of redundancies, which means that many physiological parameters can be combined into a few, non-redundant model parameters. As this manuscript will illustrate, such steps simplify both the equations and the computational handling of the model. However, in order to draw conclusions from the model calculations, the model output must be translated from mathematical language to biology. The challenge in understanding and the art of physiological interpretation lies particularly in the fact that independent physiological parameters in biology can be redundant in mathematics. Having this in mind, we may now tackle the question:

How can the net-fluxes through a membrane transporter in steady state be described?

Steady state corresponds to the state in which the system does not change any longer. Indeed, homeostatic conditions are steady state conditions because the system exhibits a certain stability and maintains its parameters constant. Homeostatic properties of a system can be analyzed in a two-step process. First, the steady state is analyzed and then, in a second step, the system is destabilized in order to investigate its resilience. In this study, we begin with the first step and will present the second in a follow-up paper.

At the beginning, we consider passive or secondary active transporters, i.e. membrane proteins that do not couple ATP hydrolysis to transport activity (pumps will be considered later). The flux through a passive transporter (*J*_*X*_) depends on the electrochemical gradient (Δ*μ*) of the transported solute across the membrane. For a permeating substrate X, this gradient is given by:

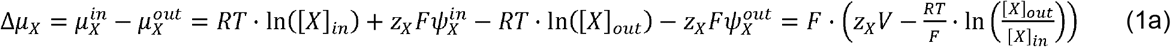

with the gas constant *R*, the Faraday constant *F*, the absolute Temperature *T*, the internal and external electrical potentials 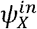 and 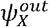 (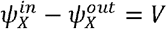, the membrane voltage), the valence *z*_*X*_ and the concentrations [*X*]_*in*_ and [*X*]_*out*_, of the substrate X, respectively (Carpaneto et al., 2005; Nour-Eldin et al., 2012). The net flux of X through the transporter (*J*_*X*_) is a function of the gradient Δ*μ*_*X*_, i.e. the concentrations and, if *Z*_*X*_≠0, also the membrane voltage. If the gradient is zero, Δ*μ* =0, there is not net flux, i.e. *J*_*X*_ = 0. Thus, if *J*_*X*_ depends on the membrane voltage, it is zero at *V*=*E*_*X*_, with 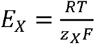.

In 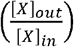, also known as the Nernst (or zero-flux) potential for X. Otherwise, it is zero if [X]_*out*_ = [X]_*in*_.

In case of a secondary active co-transporter that transports *n*_*X*_ molecules/ions of type X (valence *z*_*X*_) along with *n*_*Y*_ molecules/ions of type Y (valence *z*_*Y*_), the electrochemical gradient is given by:

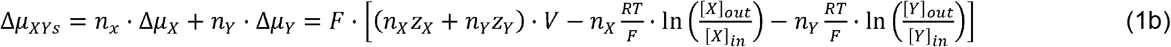

for a symporter, and by

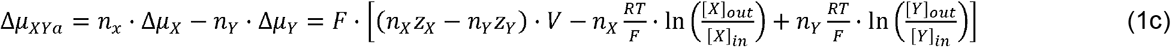

for an antiporter. In case of symporters, the flux *J*_*XY*_ is zero at 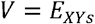, with 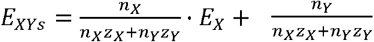. In case of antiporters, it is zero at 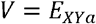, with 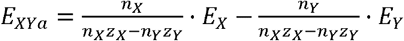, if *n*_*X*_*Z*_*X*_ − *n*_*Y*_*Z*_*Y*_ *≠* 0; otherwise it is zero if *n*_*X*_ · *E*_*X*_ = *n*_*Y*_ · *E*_*Y*_

As we will show now, this rudimentary information on Δ*μ* = 0 and *J* = 0 can be used to mathematically describe the flux in the steady state even without further knowledge of the function *J*_*X*_(*V*). The function *J*_*X*_(*V*) describes a curve in a 2-dimensional space (**Figure 2A**). We know from above consideration that *J*_*X*_(*E*_*X*_) = 0 (**Figure 2**, *blue point*). This point is strategic because at *E*_*X*_ the direction of the flux changes, e.g. if *J*_*X*_ < 0 at *V* < *E*_*X*_, then *J*_*X*_ > 0 at *V* > *E*_*X*_ (**Figure 2**, *grey regions*). In the steady state, both the membrane voltage and the flux through the transporter are constant. Thus, in this case we need to describe just one point of the *J*_X_(*V*) curve (**Figure 2B**, *red point*). For such a description two coordinates are sufficient: One is the membrane voltage in steady state, *V*_*ss*_, and the other is the flux in steady state,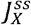. Using triangulation, 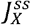 can be replaced by (**Figure 2B**, *orange triangle*)

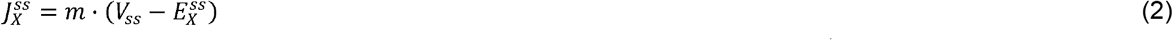

with slope *m* of magnitude *m* = tan (*α*). Flux values, *J*_*X*_(*V*) have the unit s^−1^, while membrane voltage and *E*_*X*_ have the unit V. Therefore, the slope *m* has the unit V^−1^·s^−1^. We can consider it as a free parameter (ranging from zero to infinity), which allows to reach any point on the vertical green line in **Figure 2B**. In this way, any *J*_*X*_(*V*) curve is represented by 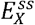, *V*_*ss*_ and the parameter *m*, even without knowing the exact shape of *J*_*X*_(*V*). *m* combines several physiologically important parameters because a change in the activity of the transporter protein is reflected by a change of the slope *m* (i.e. the angle *α*). An increase in activity, for instance by enhanced expression of the transporter or its activation by posttranslational modifications, results in higher *J*-values (**Figure 2C**, *dashed grey line*, purple point). This is reflected by a larger angle (*α*1) and a larger slope (*m1*). Conversely, a reduction in activity (**Figure 2C**, *dotted grey line, pink point*) results in smaller *J*-values, a smaller angle (*α*2) and a smaller slope (*m2*). Thus, the parameter *m* gathers all regulatory features of the respective transporter. It can take on any value in the interval [0,∞). The rather simple mathematical description of eqn. (2) therefore covers the entire range of possibilities without any constraints of approximations.

**Figure 2.**
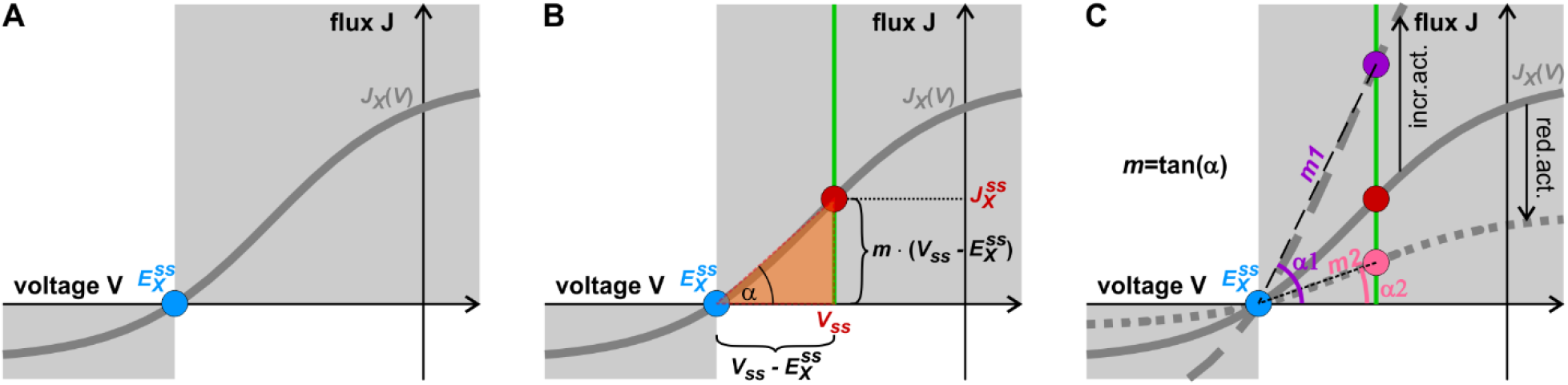
Modelling of fluxes through a transporter in steady state. (**A**) The flux curve of a transporter *J*_*X*_(*V*) (*gray line*) is zero at 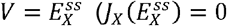, *blue point*). (**B**) In steady state, flux and voltage are constant, and the curve is represented by the *red point* (*V*_*ss*_ | *J*_*X,ss*_). The abscissa of this point is *V*_*ss*_, while the ordinate, 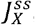, can be expressed as 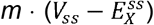 using triangulation (*m* = tan(*α*)). (**C**) Changes in the activity are mirrored by changes in the slope *m* and the angle *α*. An increase in the transporter activity (*dashed grey line*), e.g. by a higher expression or activation by phosphorylation, results in a larger 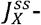 value (*purple point*), which is represented by a larger angle (*α*1) and hence a larger slope (*m1*). A decrease in transporter activity (*dotted grey line*), on the other hand, results in a smaller 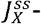value (*pink point*), smaller angle (*α*2) and smaller slope (*m2*). Thus, the slope *m* contains the regulatory features of the transporter.

To illustrate this approach in practical terms, we develop it in the following, step by step, for (i) K^+^ channels, (ii) H^+^/K^+^ symporters, and (iii) H^+^/K^+^ antiporters. Finally, we will also introduce a suitable model for the (iv) H^+^-ATPase that primarily energizes all transport processes.

#### (i) K^+^ channels

The electrochemical gradient (compare eq. 1a) that determines the flux across K^+^ channels is given in the case of highly selective K^+^ channels by:

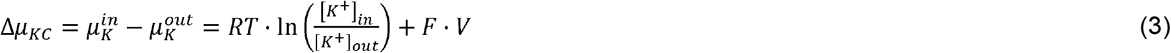

with the internal and external K^+^ concentrations [*K*^*+*^]_*in*_ and [*K*^*+*^]_*out*_, respectively. This gradient is zero at the zero-flux potential for K^+^ channels, *V* = *E*_*K*_, which is the Nernst potential for potassium

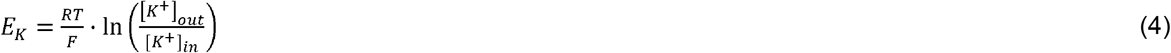

Thus, at the *V* = *E*_*K*_ function of the flux, *J*_*K,KC*_(*V*)(unit s^−1^), has the value *J*_*K,KC*_(*E*_*K*_) = 0. Applying the same considerations that led to equation (2), the flux in steady state through the K^+^ channel can be described by:

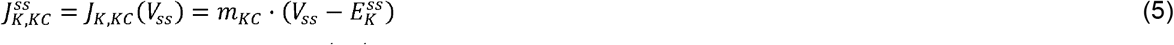

The parameter *m*_*KC*_ (unit V^−1^·s^−1^) determines the activity of the channels and comprises all potential regulatory features, such as functional protein expression, post-translational modifications and/or voltage dependence. The net K^+^ flux through the K^+^ channels causes a transmembrane electric current (unit A) given by:

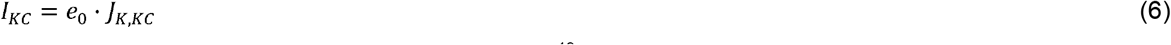

with the elementary (positive) charge *e*_*0*_ (= 1.6×10^−19^ A·s).

#### (ii) H^+^/K^+^ symporters

The electrochemical gradient (compare eq. 1b) that determines the flux across a symporter with *n*_*S*_ H^+^ / 1 K^+^ stoichiometry is given by:

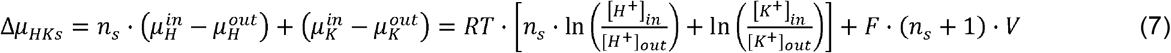

with the internal and external proton concentrations [*H*^*+*^]_*in*_ and [*H*^*+*^]_*out*_, respectively. This gradient is zero at the zero-flux potential *V* = *E*_*HKs*_ with

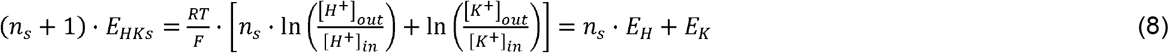

and the Nernst potentials for potassium (*E*_*K*_) and protons (*E*_*H*_).

Hence, the net-fluxes for K^+^ and protons, respectively, at *V* = *E*_*HKs*_ are *J*_*K,HKs*_(*E*_*HKs*_) =0 and *J*_*H,HKs*_(*E*_*HKs*_) = *n*_*s*_· *J*_*K,HKs*_(*E*_*HKs*_) = 0. Applying the same considerations that led to equation (2), the fluxes in steady state can be described by:

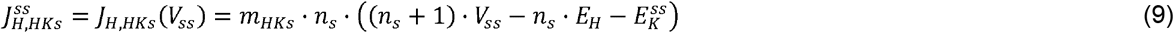

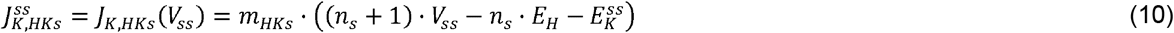

Also here, the parameter *m*_*HKs*_ gathers all regulatory features of the symporter. The transmembrane electric current (unit A) that is caused by the net H^+^ and K^+^ fluxes (unit s^−1^) through the H^+^/K^+^ symporters is given by:

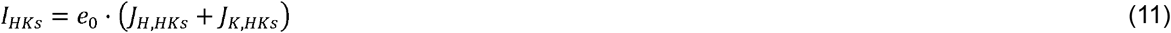

#### (iii) H^+^/K^+^ antiporters

The electrochemical gradient (compare eq. 1c) that determines the flux across an antiporter with *n*_*a*_ H^+^ / 1 K^+^ stoichiometry is given by:

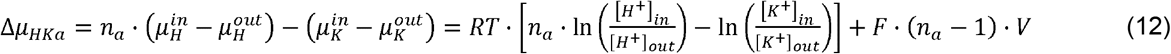

This gradient is zero at *E*_*HKa*_ with

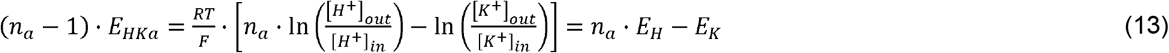

If we apply again the same considerations that led to equation (2), the net-fluxes for K^+^ and H^+^ in steady state can be expressed as:

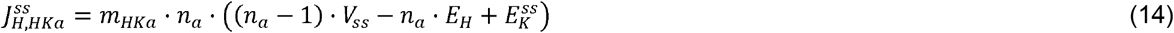

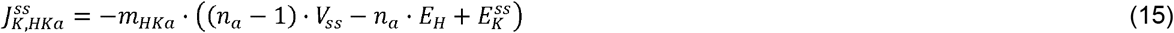

Here as well, the parameter *m*_*HKa*_ combines all regulatory characteristics of the antiporter. The transmembrane electric current (unit A) that is caused by the net H^+^ and K^+^ fluxes (unit s^−1^) through the H^+^/K^+^ antiporters is given by:

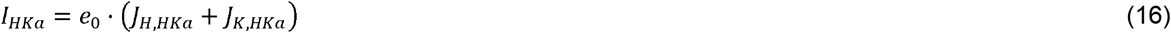

It is zero, if *n*_*a*_ = 1. In this case, the antiport would be electroneutral and the flux would not depend on the membrane voltage, but would only be proportional to the difference *E*_*H*_ − *E*_*K*_ between the Nernst potentials of H^+^ and K^+^.

#### (iv) H^+^-ATPase

Finally, we consider the proton pump as active membrane transporter. The derivation of a mathematical description of the pump current of H^+^-ATPases was presented in detail in the Supplementary Material by Reyer et al. (2020). The mechanistic model of a pump cycle (Dreyer, 2017; Reyer et al., 2020) results in a sigmoidal curve for pump currents (unit A) that describes very well the experimental findings (Lohse and Hedrich, 1992):

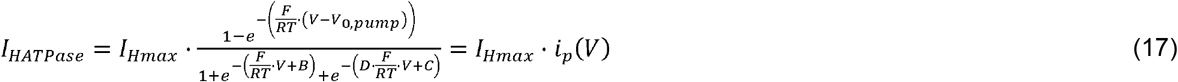

with 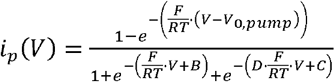. The proton flux mediated by the H^+^-ATPase (*J*_*HATPase*_; unit s^−1^) can be determined by dividing the current by the elementary (positive) charge *e*_*0*_:

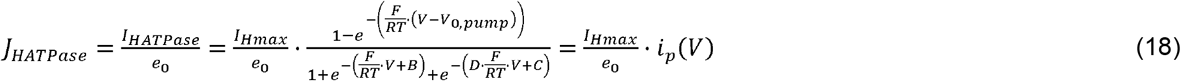

Here, the numerical values of B, C, and D are of minor importance for the shape of the curve. Suitable values are B = 5.4, C = 2.5, D = 0.1 (Dreyer, 2021a). Most important are the parameters *I*_*Hmax*_ and *V*_*0,pump*_. *I*_*Hmax*_ determines the maximum pump current and is proportional to the number of active proton pumps in the membrane, while *V*_*0,pump*_ is the voltage at which the pump current is zero [*i*_*p*_(*V*_*0,pump*_) = 0]. It depends in a complex manner on the cytosolic and apoplastic proton concentrations and on the cytosolic ATP, ADP, and Pi concentrations (Rienmüller et al., 2012). For suitable simulations, in which the energy status of the cell does not change in the considered time interval, *V*_*0,pump*_ can be set and kept constant (e.g. *V*_*0,pump*_ = −200 mV). This value determines the most negative value that the membrane voltage can attain.

### Mathematical Description of changes in concentrations and voltage

The proton and K^+^ fluxes change the concentrations of H^+^ and K^+^ on both sides of the membrane. The combined net K^+^ efflux (*J*_*K*_ = *J*_*K,KC*_ + *J*_*K,KHs*_ + *J*_*K,KHa*_; unit s^−1^) increases [*K*^*+*^]_*out*_ and decreases [*K*^*+*^]_*in*_, while the combined H^+^ efflux (*J*_*H*_ = *J*_*HATPase*_ + *J*_*H,KHs*_ + *J*_*H,KHa*_; unit s^−1^) increases [*H*^*+*^]_*out*_ and decreases [*H*^*+*^]_*in*_. Additionally, the buffer capacities of the internal and external compartments can partially mitigate the changes in proton concentration, described by the general buffer reaction 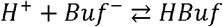 with an anionic base *Buf*^*−*^ and its conjugate acid, *HBuf*. All these changes are represented in mathematical terms by ordinary differential equations. The changes in K^+^ concentrations are governed by:

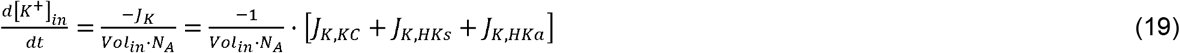

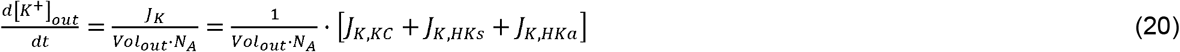

Here *N*_*A*_ is the Avogadro constant and *Vol*_*in*_ and *Vol*_*out*_ are the volumes of the compartments on the respective side of the membrane. Changes in proton concentrations are determined by:

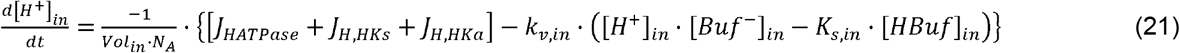

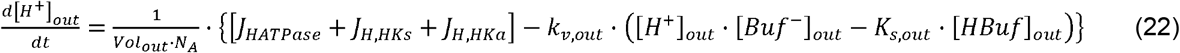

Here, 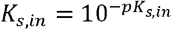 and 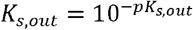 indicate the dissociation constants of the buffer reactions that buffer *pH*_*in*_ and *pH*_*out*_ to *pK*_*s,in*_ and *pK*_*s,out*_, respectively. The buffer concentrations, in turn, change according to:

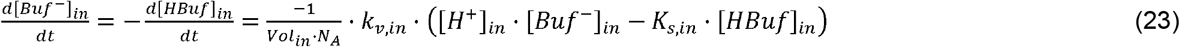

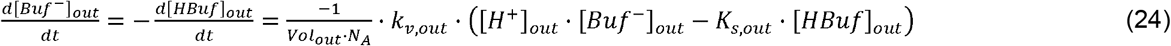

Compared to the transport reactions, the buffering reactions take place very quickly, which is reflected in large *k*_*v,in*_ and *k*_*v,out*_ values. Because of this difference in time-scale, the parameters *k*_*v,in*_ and *k*_*v,out*_ do not need to be known in detail. In simulations, it is sufficient to set them 2-3 magnitudes larger than the other parameters (e.g. *k*_*v,[in,out]*_/(*I*_*max*_/*e*_*0*_) = 100 *μ*M^−2^) to obtain almost instantaneous buffer reactions in comparison to the time-scale of transport.

The net fluxes usually cause a net charge transport across the membrane. This non-zero current provokes a change of the membrane voltage. The membrane is actually comparable to a dielectric in an electric capacitor that separates the charges in the aqueous interior from the aqueous exterior. Charge transport from one side to the other changes the membrane voltage according to:

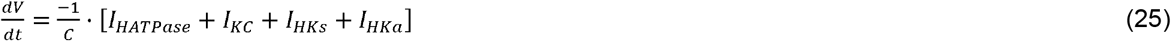

with the membrane capacitance *C* (unit F).

### Modeling the steady state with fixed transporters activities

At this point, we have translated figure 1 into the language of mathematics. For each transporter, we expressed the fluxes and currents in equations (5, 6, 9, 10, 11, 14, 15, 16, 17, 18) with a few parameters. The dynamic properties of the system are determined by equations (19-25). In the following, we will solve the differential equations for homeostatic conditions. If the system is considered in steady state, neither the membrane voltage nor the concentrations change with time, which means that the left sides of equations (19-25) are zero. In this condition, the equations (19-25) simplify to two non-redundant equations taking also into account that *I*_*X*_ = *e*_*0*_·*J*_*X*_:

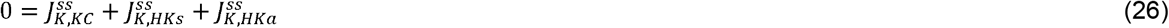

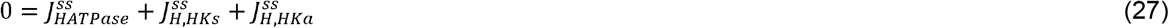

All the other equations are either zero or linear combinations of the equations (26-27). In the next step, the 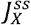 were replaced by the expressions deduced in equations (5, 9, 10, 14, 15, 18) yielding:

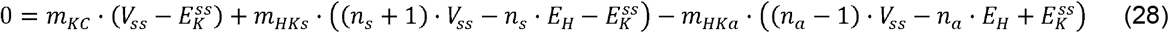

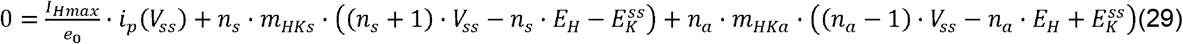

Then, we multiplied both equations with the factor *e*0/*I*_*Hmax*_ and define *g*_*X*_ ≔ *m*_*x*_*· e*_0_/*I*_*Hmax*_ (unit V^−1^). This operation eliminated a redundant parameter and reduced the set of four parameters (*I*_*Hmax*_, *m*_*KC*_, *m*_*HKs*_, *m*_*HKa*_) to a set of three (*g*_*KC*_, *g*_*KHs*_, *g*_*KHa*_) representing the *relative* activity of the transporter proteins (relative to the maximal activity of the H^+^-ATPase).

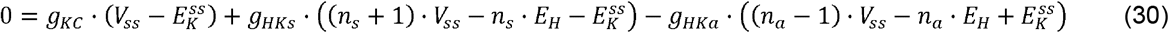

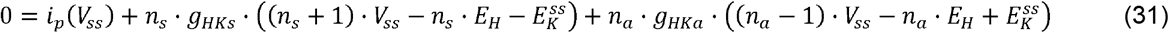

Additionally, the equations (30) and (31) could be arranged in vector and matrix form:

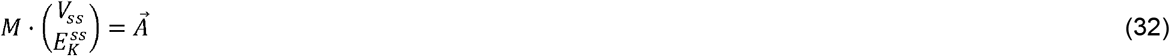

with:

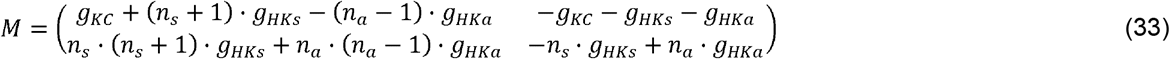

and:

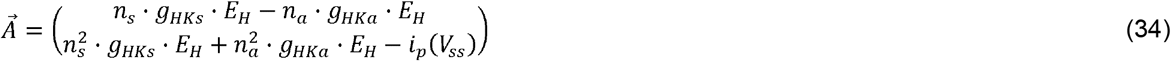

This might be considered as just a formalism and not needed for two equations. However, the conversion should illustrate that we have opened the powerful toolbox of linear algebra. More complex homeostats with more than two equations can be handled in the same way and the tools from linear algebra for solving this type of equation can be applied in the same manner. Equation (32) has exactly one solution for the 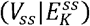 pair that depends on the activities of the transporter proteins (*g*_*X*_) and the proton gradient (*E*_*H*_):

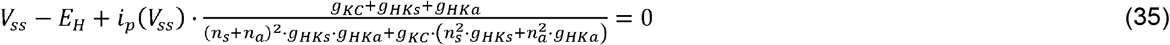

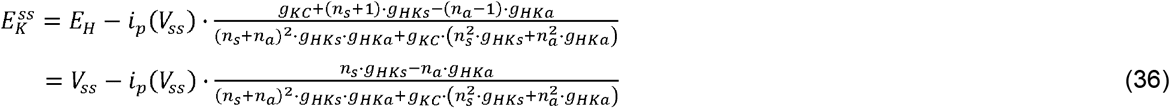

Please note

1. Equations (35, 36) are not defined for cases in which two of the three parameters *g*_*KC*_, *g*_*HKs*_, *g*_*HKa*_ are zero. In cases, in which only one *g*_*X*_ is different from zero, the equations (30, 31) result in:

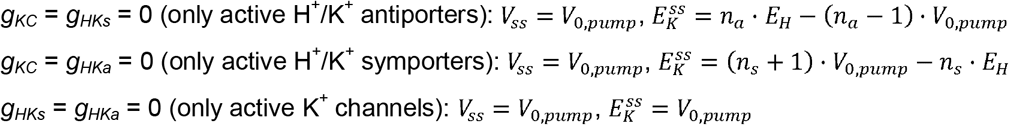
2. Equation (35; of type *V*_*ss*_ − *A* +*i*_*p*_ *V*_*ss*_) ·*B*= 0) is an implicit solution. It cannot be explicitly solved analytically for *V*_*ss*_ because *i*_*p*_(*V*_*ss*_) is a function of *V*_*ss*_. Nevertheless, mathematical tools also allow to bypass this difficulty. The left-hand side of eqn. (35) before the equals sign is a strictly monotonically increasing function of *V*_*ss*_, which means that the first derivative after *V*_*SS*_ is > 0. Or in other words, with increasing *V*_*ss*_, the left-hand side of the equation also increases. The corresponding curve therefore cuts the zero axis only once. Thus, for given values of *A* =*E*_*H*_ and *B*= *f*(*g*_*KC*,_*g*_*HKs*,_*g*_*HKa*_) there is exactly one *V*_*ss*_ that obeys this equation. If *E*_*H*_, *g*_*KC*_, *g*_*HKs*_, and *g*_*HKa*_ are known, the value of *V*_*ss*_ can be determined numerically by root-finding algorithms, such as the Newton–Raphson method.

## Results

The presented thermodynamical and mathematical analysis of the transporter network has resulted in general solutions for (eqn. 35) and 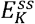 (eqn. 36) in the steady state, i.e. in homeostatic conditions. These values could be influenced by the cell via the proton gradient *E*_*H*_ and the parameters *g*_*KC*_, *g*_*HKs*_, and *g*_*HKa*_, i.e. the (relative) activities of the different K^+^ transporters. Considering the fact that from a physiological point of view the proton gradient is less flexible, the main setscrews to adjust 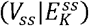 are the *g*_*X*_ values. These three mathematical parameters split in the biological reality into a manifold of physiological parameters, such as gene expression, protein turnover and post-translational regulation. Thus, one *g*_*X*_ value can represent several different biological realities.

To illustrate the consequences of the dependency of the homeostatic steady state on the parameters *g*_*KC*_, *g*_*HKs*_, and *g*_*HKa*_, the results for the case *n*_*s*_ = 1, *n*_*a*_ = 1, *V*_*0,pump*_ = −200 mV, and *E*_*H*_ = +57.6 mV (pH_in_ = 7, pH_out_ = 6) are shown as an example. It should be noted that another choice of these parameters would not have changed the results qualitatively. For each parameter *g*_*X*_, we chose 0 and 19 logarithmically distributed values in a range between 10^−6^ mV^−1^ and 1 mV^−1^ (i.e. 0 mV^−1^, 1×10^−6^ mV^−1^, 2×10^−6^ mV^−1^, 5×10^−6^ mV^−1^, 1×10^−5^ mV^−1^, 2×10^−5^ mV^−1^, 5×10^−5^ mV^−1^, 1×10^−4^ mV^−1^, 2×10^−4^ mV^−1^, 5×10^−4^ mV^−1^, 1×10^−3^ mV^−1^, 2×10^−3^ mV^−1^, 5×10^−3^ mV^−1^, 1×10^−2^ mV^−1^, 2×10^−2^ mV^−1^, 5×10^−2^ mV^−1^, 1×10^−1^ mV^−1^, 2×10^−1^ mV^−1^, 5×10^−1^ mV^−1^, 1 mV^−1^). With respect to **Figure 2B**, this range covered a spectrum of the angle *α* between 0° and almost 90° and thus the entire range of possibilities. The resulting 20×20×20 table represented 8000 different (mathematical) realities of the system. For each of them, eqn. (35) was solved using the “Goal Seek” routine in Excel, setting eqn. (35) to 0 by changing *V*_*ss*_. This was done to show that no sophisticated computer programs are required for the analyses. With the resulting value for, then the value for 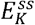 was calculated using eqn. (36). The determined 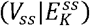 pair finally allowed to calculated the different relative fluxes in steady state using eqns. (5, 9, 10, 14, 15, 18) and the definition *g*_*X*_ ≔ *m*_*x*_*· e*_0_/*I*_*Hmax*_ (see above):

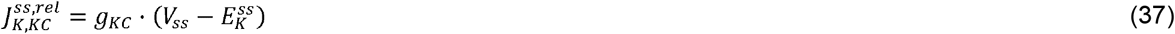

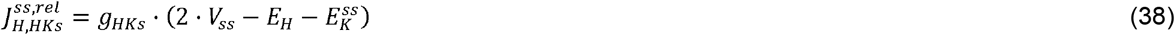

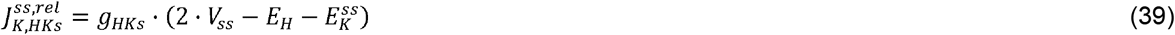

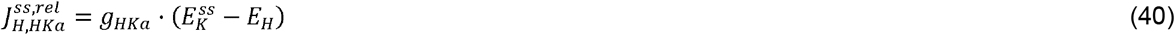

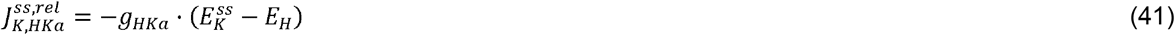

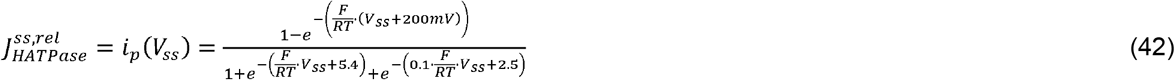

### K^+^ and H^+^ cycling in steady state

In the 8000 different cases, *V*_*ss*_ ranged between −200 mV (= *V*_*0,pump*_) and +57.6 mV (= *E*_*H*_), while 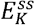 assumed values between −457.6 mV (= 2· *V*_*0,pump*_ − *E*_*H*_) and +57.6 mV (= *E*_*H*_). The **Figure 3** illustrates the values of *V*_*ss*_ (A,D,G),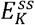 (B,E,H), and 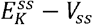 (C,F,I) for several parameter sets of *g*_*KC*_, *g*_*HKs*_, and *g*_*HKa*_. The larger was the activity of the K^+^ channel (B<E<H), the less negative was the minimal 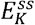 that could be achieved by highly active H^+^/K^+^ symporters. With hardly any active K^+^ channels (*g*_*KC*_ = 10^−5^ V^−1^) the symporter could (theoretically) accumulate a [K^+^]_in_ that was almost 10^8^-fold higher than [K^+^]_out_, which corresponded to 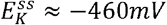 (**Figure 3B**). In this condition, the membrane voltage was *V*_*ss*_ *≈ −*200 *mV* (**Figure 3A**), i.e. close to the limit of the pump *V*_*0,pump*_. This means that all energy from the proton gradient and ATP-hydrolysis was used for K^+^ accumulation. These extreme values became more moderate with increasing channel activity. At 500-fold higher K^+^ channel activity (*g*_*KC*_ = 5×10^−3^ V^−1^), 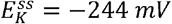 was the most negative value at high *g*_*HKs*_ and low *g*_*HKa*_ (**Figure 3E**), which still corresponded to ~17.000-fold higher [K^+^]_in_ than [K^+^]_out_. The reduction in the accumulation rate came along with an increased *V*_*ss*_, in particular at higher symporter but also at higher antiporter activities (**Figure 3D**) pointing to an energy-dissipation process that increased with increasing K^+^ channel activity (**Figure 3A**,**D**,**G**). This was further corroborated when analyzing the transmembrane fluxes (**Figure S1**). Although the net fluxes of K^+^ and H^+^ were zero in steady state (eqns. 26-27), there was a transmembrane cycling of both ions. Permanent effluxes were compensated by permanent influxes. To illustrate the cycling, the H^+^ and K^+^ effluxes and the ratio between the two were displayed in **Figure 4** relative to the maximally achievable proton pump-driven H^+^ efflux; *i*.*e*. the maximum H^+^ efflux of the pump (*I*_*Hmax*_/*e*_*0*_, eqn. 18) was normalized to 1. If the K^+^ channel activity was very low (*g*_*KC*_ = 10^−5^ V^−1^), the relative H^+^ efflux mediated by the pump ranged from 0 to ≈0.94 [=*i*_*p*_(*E*_*H*_)] with the highest value achieved with highly active sym- and antiporters (**Figures 4A, S1A**) at *V*_*ss*_ = *E*_*H*_ (**Figure 3A**). Under these conditions, the relative K^+^ efflux was half as large as the H^+^ efflux (**Figure 4B**), indicating that two protons were looped for one K^+^ ion (**Figure 4C**). And indeed, one H^+^ was taken up together with one K^+^ by the H^+^/K^+^ symporter (**Figure S1B**,**E**), while the other H^+^ was taken up by the H^+^/K^+^ antiporter releasing one K^+^ (**Figure S1C**,**F**). The accumulated protons were released by the pump (**Figure S1A**)

**Figure 3.**
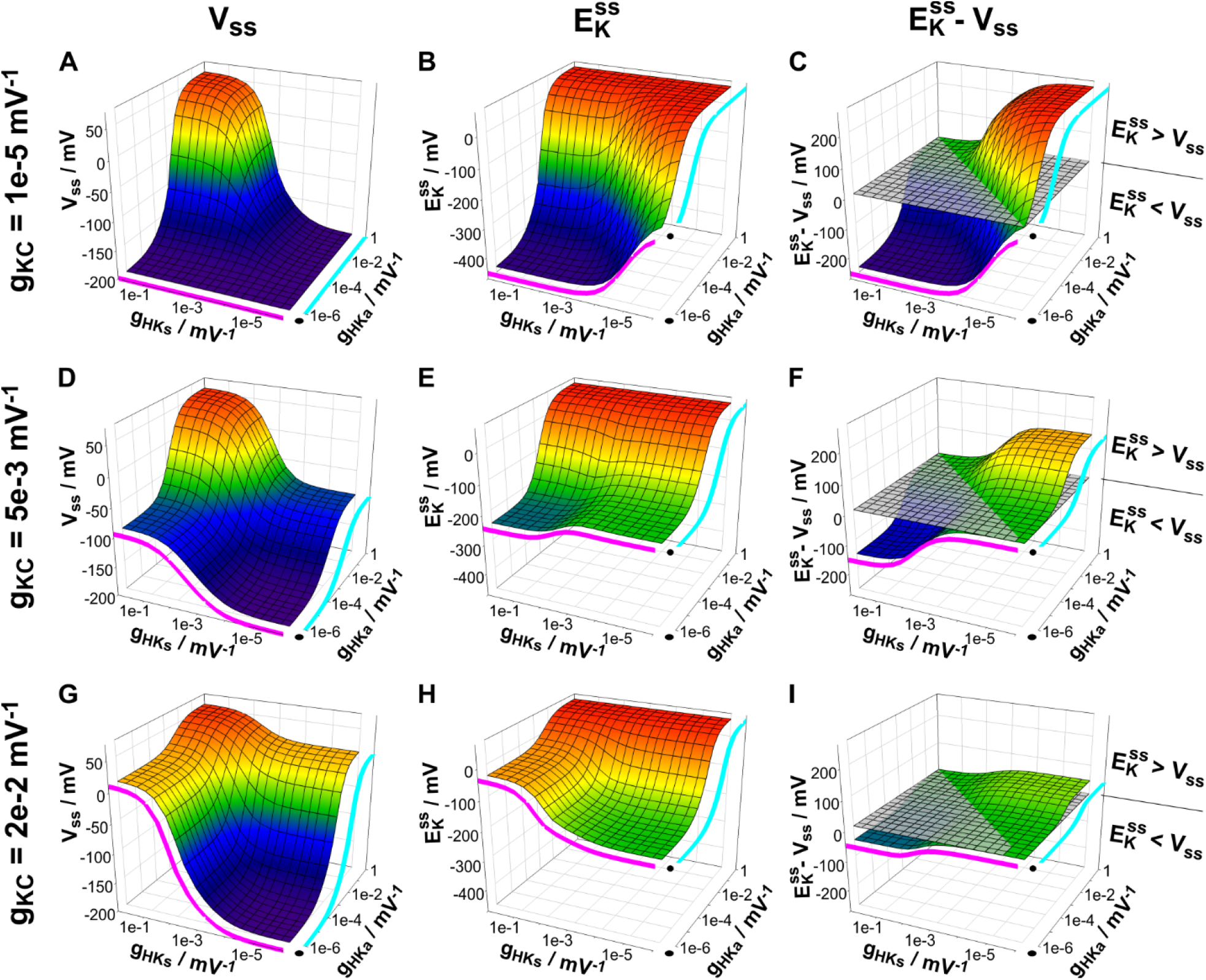
Membrane voltage and K^+^ gradient in homeostatic (steady state) conditions. The activities of K^+^ channels (*g*_*KC*_), H^+^/K^+^ symporters (*g*_*HKs*_) and H^+^/K^+^ antiporters (*g*_*HKa*_) determine the membrane voltage (*V*_*ss*_, **A, D, G**) and the K^+^ gradient (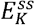, **B, E, H**) in steady state. The difference 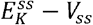 (**C, F, I**) is positive if the activity of antiporters is higher than that of symporters (*g*_*HKa*_ > *g*_*HKs*_). It is negative if *g*_*HKa*_ > *g*_*HKs*_. Data were calculated for the case *n*_*s*_ = 1, *n*_*a*_ = 1, *V*_*0,pump*_ = −200 mV, and *E*_*H*_ = +57.6 mV (ΔpH = 1). The magenta lines show the values in the absence of active H^+^/K^+^ antiporters (*g*_*HKa*_ = 0), whereas the cyan lines indicate the values in the absence of active H^+^/K^+^ symporters (*g*_*HKs*_ = 0).

**Figure 4.**
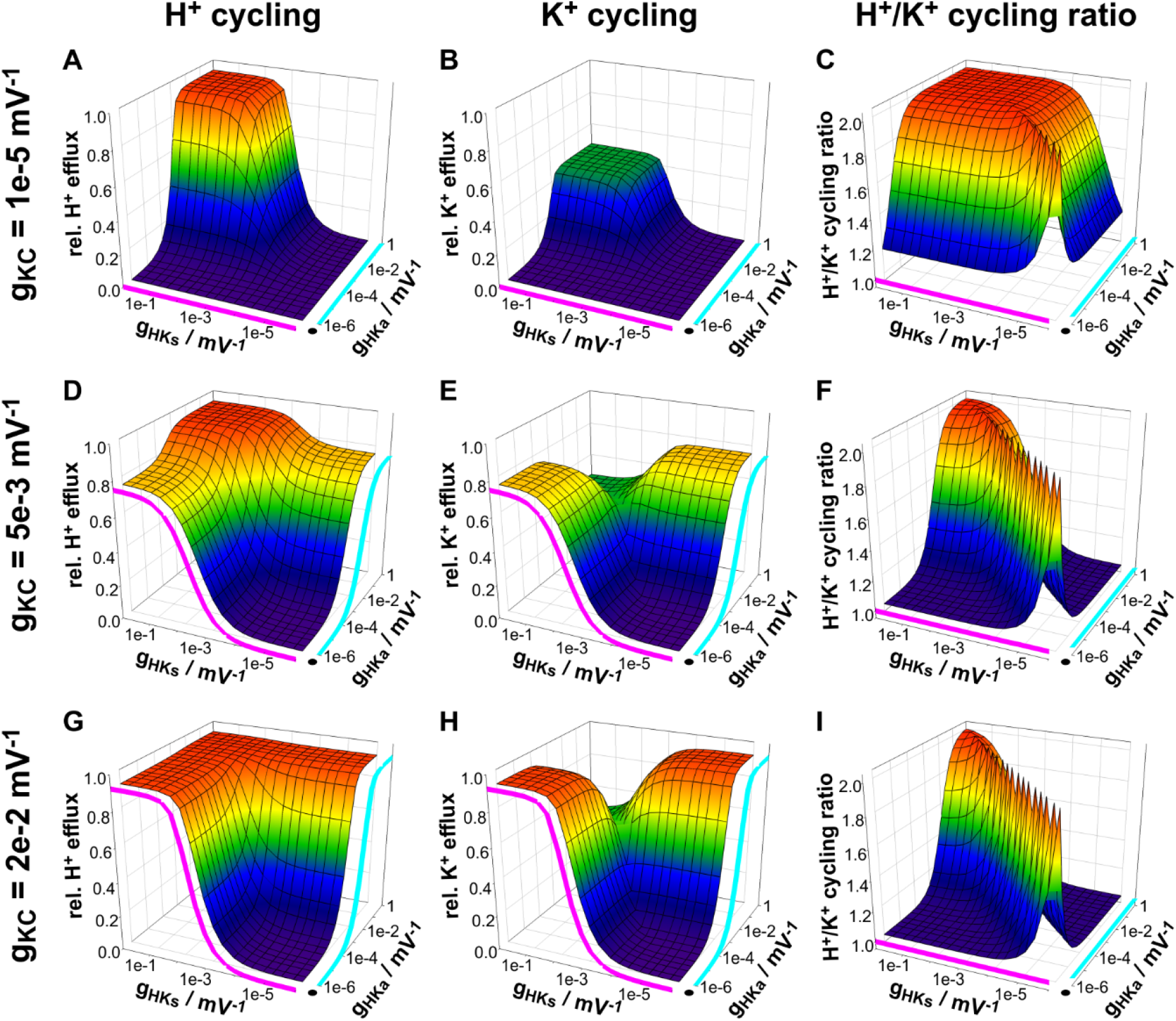
H^+^ and K^+^ cycling in homeostatic (steady state) conditions. Although the net H^+^ and K^+^ fluxes are zero in homeostatic conditions, there are still continuous H^+^ and K^+^ effluxes that are compensated by influxes of the same magnitude. (**A, D, G**) Dependency of the H^+^ efflux and (**B, E, H**) of the K^+^ efflux on the activities of the K^+^ channels (*g*_*KC*_), H^+^/K^+^ symporters (*g*_*HKs*_) and H^+^/K^+^ antiporters (*g*_*HKa*_). The effluxes are shown relative to the maximal H^+^ efflux that can be generated by the H^+^ ATPase (*J*_*Hmax*_ = *I*_*Hmax*_/*e*_*0*_). (**C, F, I**) Ratio between H^+^ efflux and K^+^ efflux as a measure for the H^+^/K^+^ cycling ratio. Data were calculated for the case *n*_*s*_ = 1, *n*_*a*_ = 1, *V*_*0,pump*_ = −200 mV, and *E*_*H*_ = +57.6 mV (ΔpH = 1). The magenta lines show the values in the absence of active H^+^/K^+^ antiporters (*g*_*HKa*_ = 0), whereas the cyan lines indicate the values in the absence of active H^+^/K^+^ symporters (*g*_*HKs*_ = 0).

H^+^ and K^+^ effluxes became smaller with either decreasing symporter or decreasing antiporter activity (**Figure 4A**,**B**). This picture changed with increasing K^+^ channel activity. While the relative H^+^ and K^+^ effluxes remained unaffected by an increased channel activity at very high sym- and antiporter activity (compare the plateaus in the corners of **Figures 4A**,**B**,**D**,**E**,**G**,**H**), K^+^ channel activity had a strong effect when one of the H^+^/K^+^ transporters was less active. With less active antiporters and highly active symporters, increased K^+^ channel activity increased the K^+^ efflux (compare **Figures 4B**,**E**,**H**, close to the magenta curve). This K^+^ efflux was mediated by the K^+^ channels (**Figure S1D**,**J**,**P**), compensated by a larger K^+^ influx via the H^+^/K^+^ symporters (**Figure S1E**,**K**,**Q**). The accompanying H^+^ influx (**Figure S1B**,**H**,**N**) was neutralized by the H^+^ efflux via the pump (**Figure S1A**,**G**,**M**) that also energized the combined H^+^/K^+^ cycling.

Likewise, K^+^ efflux also increased with increased K^+^ channel activity at highly active antiporters and less active symporters (compare **Figures 4B**,**E**,**H**, close to the cyan curve). However, this time, the efflux was not mediated by the K^+^ channels (**Figure S1D**,**J**,**P**). They mediated K^+^ influx instead. The K^+^ efflux passed through the H^+^/K^+^ antiporters (**Figure S1F**,**L**,**R**). When comparing the two situations, it turned out that in homeostatic steady state conditions K^+^ channels served at the same membrane voltage, *V*_*ss*_, once as uptake and once as release channels depending on the H^+^/K^+^ transporter situation. In combination with H^+^/K^+^ symporters, they acted as K^+^ release channels, even at very negative voltages. In contrast, in combination with H^+^/K^+^ antiporters, they functioned as K^+^ uptake channels, even at positive voltages. These thermodynamically derived properties of K^+^ channels under homeostatic conditions obviously contradict the historically established separation of voltage-dependent K^+^ channels into hyperpolarization-activated uptake (K_in_) and depolarization-activated release (K_out_) channels (Dreyer and Blatt, 2009; Dreyer and Uozumi, 2011; Sharma et al., 2013; Dreyer et al., 2021). Thermodynamic considerations have now taught us that K_in_ channels might also serve as K^+^ release channels to avoid over-accumulation of K^+^ by H^+^/K^+^ symporters, while K_out_ channels may also serve as K^+^ uptake channels under certain homeostatic conditions.

### Conclusions on the cost efficiency of homeostasis

The considerations based on thermodynamic first principles indicated that homeostatic conditions can be achieved by combining two different K^+^ transporter types (K^+^ channels and H^+^/K^+^ symporters, **Figure 5A**, K^+^ channels and H^+^/K^+^ antiporters, **Figure 5B**, H^+^/K^+^ symporters and H^+^/K^+^ antiporters, **Figure 5C**), or all three K^+^ transporter types (**Figure 5D**). In all these cases, the homeostatic condition is inevitably accompanied by energy-consuming H^+^ and K^+^ loops across the membrane (Dreyer, 2021a). In **Figures 3, 4, and S1** this was shown exemplarily for sym- and antiporters with a 1 H^+^:1 K^+^ stoichiometry. In the following, the more general case (*n*_*s*_ and *n*_*a*_) was considered to find out which values for *n*_*s*_ and *n*_*a*_ make the most sense. The H^+^ and K^+^ cycles occurred because the homeostatic steady state was different from the equilibrium states of the different involved transporters. In the case of a transporter network with H^+^ pump, K^+^ channel and H^+^/K^+^ symporter (**Figure 5A**), for instance, the steady state membrane voltage, *V*_*ss*_, is different from the equilibrium voltages of the pump (*V*_*0,pump*_), the channel 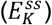 or the symporter 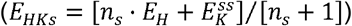. This means that the pump continuously pumps protons from the inside to the outside by consuming ATP. These protons return to the cell via the H^+^/K^+^ symporter and shuttle 1 K^+^ per *n*_*s*_ protons into the cell. The absorbed K^+^ is released again via the channel. These combined H^+^ and K^+^ cycles are yield-neutral in steady state, i.e. they do not affect neither the concentrations nor the electrical charges at both sides of the membrane. The cost of the homeostatic condition for the cell is in this case *n*_*s*_ ATP per 1 looped K^+^. Most cost-efficient would therefore be *n*_*s*_ = 1. And indeed, even at this lowest coupling rate, the H^+^/K^+^ symporter is powerful enough to (theoretically) accumulate K^+^ under physiological voltage (V ≥ −200 mV) and pH (ΔpH = 1) conditions more than 10^5^-fold in the interior compared to the exterior 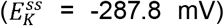. Higher coupling rates (*n*_*s*_ > 1) would allow higher accumulation rates, but at the expense of higher ATP consumption. Considering the physiological conditions, the best cost-benefit ratio is therefore *n*_*s*_ = 1. The K^+^ transporter network shown in **Figure 5A** is very well suited for K^+^ accumulation, in particular when [*K*^*+*^]_*out*_ is small. However, it has the potential problem of K^+^ overaccumulation under certain conditions (moderate [*K*^*+*^]_*out*_). The grey area in **Figure 5A** (lower panel) indicates the reachable range for *V*_*ss*_ and 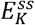. This transporter network only allows steady state conditions with 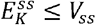, which might imply rather high [*K*^*+*^]_*in*_. This drawback could be eliminated by replacing the H^+^/K^+^ symporter by an H^+^/K^+^ antiporter (**Figure 5B**). The network of pump, channel and antiporter can establish homeostatic conditions with 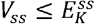, and might therefore be suitable for moderate [*K*^*+*^]_*out*_. In steady state, the protons pumped out of the cell are reaccumulated by the antiporter, which releases K^+^, that is then reabsorbed by the channel. The cost of these H^+^ and K^+^ cycles is *n*_*a*_ ATP per 1 looped K^+^. The *n*_*a*_ value only affects the upper limit of 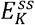. For 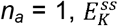 cannot be larger than *E*_*H*_ (**Figure 5B**, lower panel, grey area). Higher *n*_*a*_ values would allow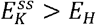. However, this is physiologically not relevant because it would mean very low [*K*^*+*^]_*in*_. Thus, H^+^/K^+^ antiporters with *n*_*a*_ =1 are the most cost-efficient, i.e. they function as electroneutral transporters. The limits of the two formerly considered networks can be overcome by combining the H^+^ pump with H^+^/K^+^ sym- and antiporters (**Figure 5C**). Here, the combined range of 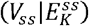 pairs is in reach (*light* and *dark grey areas*). The drawback is, however, the increased cost of (*n*_*s*_+*n*_*a*_) ATP per 1 looped K^+^, i.e. 2 ATP per 1 cycled K^+^ in the most cost-efficient case. The cost can be reduced by including additionally K^+^ channels into the transporter network (**Figure 5D**). Without any compromise on the range of the reachable 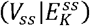 pairs, 1 looped K^+^ costs here 1-2 ATP (in the most cost-efficient case *n*_*s*_ = *n*_*a*_ = 1, **Figure 4C,F,I**).

**Figure 5.**
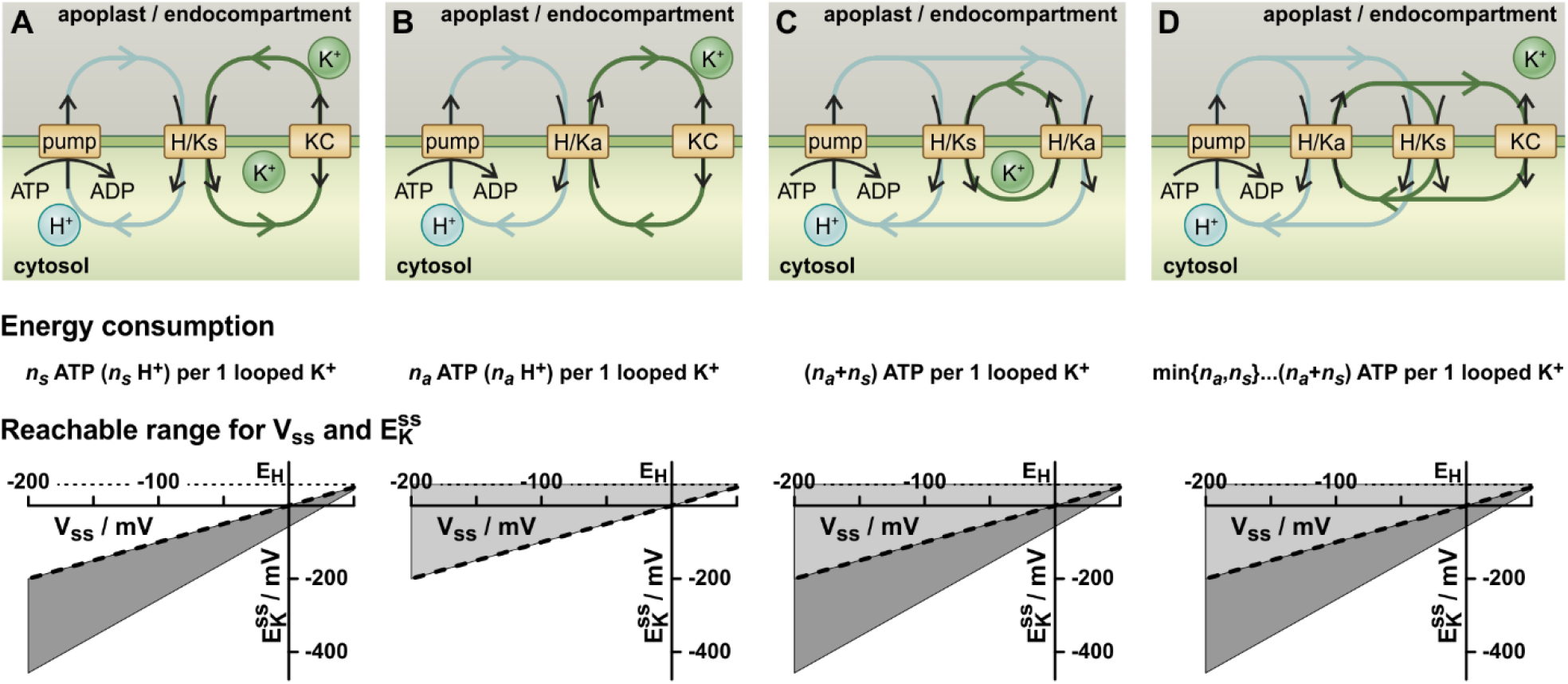
K homeostats in steady state. The three different K^+^ transporter types (i) K^+^ channels (KC), (ii) H^+^/K^+^ symporters (H/Ks, stoichiometry *n*_*s*_ H^+^ : 1 K^+^) and (iii) H^+^/K^+^ antiporters (H/Ka, stoichiometry *n*_*a*_ H^+^ : 1 K^+^) can be arranged to a K homeostat in four different combinations. In all combinations the transmembrane, yield-neutral, but energy-consuming cycling of H^+^ and K^+^ is a feature of the steady state condition. **(A)** A network of H^+^-pumps, K^+^ channels and H^+^/K^+^ symporters consumes *n*_*s*_ ATP per 1 looped K^+^, but allows only conditions for which 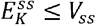 (*dark grey area*, shown for the most cost-efficient case *n*_*s*_ = 1). **(B)** A network of H^+^-pumps, K^+^ channels and H^+^/K^+^ antiporters consumes *n*_*a*_ ATP per 1 looped K^+^, but allows only conditions for which 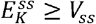 (*light grey area*, shown for the most cost-efficient case *n*_*a*_ = 1). **(C)** A network of H^+^-pumps, H^+^/K^+^ symporters and H^+^/K^+^ antiporters consumes (*n*_*s*_+*n*_*a*_) ATP per 1 looped K^+^. It allows a broader range of 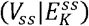 pairs (combined *grey areas*, shown for the most cost-efficient case *n*_*s*_ = *n*_*a*_ = 1). **(D)** A network of H^+^-pumps, K^+^ channels, H^+^/K^+^ symporters and H^+^/K^+^ antiporters allows the same broad range of 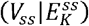 pairs as in (C) (combined *grey areas*, shown for the most cost-efficient case *n*_*s*_ = *n*_*a*_ = 1). However, the ATP consumption is smaller and ranges between 1 and 2 ATP per 1 looped K^+^ (in the most cost-efficient case *n*_*s*_ = *n*_*a*_ = 1). [Adapted from Dreyer (2021a)].

## Discussion and Conclusion

Cellular ion homeostasis is governed by fundamental physical and chemical laws. In this study, we used (i) the definition of the electrochemical potential, (ii) the basic consequence from this definition that net fluxes are zero if there is no gradient in the electrochemical potential, (iii) the chemical definition of concentrations and concentration changes, (iv) the physical properties of membranes to separate charges like a technical capacitor, and we (v) assumed that the considered transporters function as perfect molecular machines as shown exemplarily for several of them (e.g. Carpaneto et al., 2005). We described the different processes with mathematical equations and took advantage from the powerful toolboxes of differential equations and linear algebra. These basic considerations were sufficient to draw fundamental conclusions about the behavior of K^+^ transporter networks responsible for setting homeostatic (steady state) conditions for potassium in a cellular or organelle environment:

1. By setting the transporter activities (*g*_*X*_), the cell can precisely adjust the steady state values for the membrane voltage *V*_*ss*_ and the transmembrane K^+^ gradient 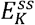. A cell can change the parameters *g*_*X*_ by altering the activities through gene expression and post-translational modifications (e.g. phosphorylation), which points to the adjusting screws that integrate the homeostats into the cellular regulatory networks.
2. The cell cannot adjust [*K*^*+*^]_*in*_ independent of [*K*^*+*^]_*out*_; it can only adjust 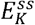, which depends on both.
3. If the membrane has only one functional K^+^ transporter type, i.e. all *g*_*X*_ except one are zero, the steady state is independent of the activity of this transporter and cannot be adjusted. In other words: Homeostatic conditions involve always at least two different types of transporters.
4. Transporter types are only different, if they are energized differently, i.e. the driving gradient Δ*μ* is different. For instance, voltage-dependent and voltage-independent K^+^ channels are not different; both are described by the same type of equations. The voltage-dependence is covered by the *g*_*KC*_ parameters, and several *g*_*KC*_ values could be combined to one that represents all channels.
5. A steady state condition of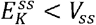, is (indirect) proof of functional H^+^/K^+^ symporters in the membrane. The condition 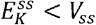 is only achievable with *g*_*HKs*_ > 0.
6. Most cost efficient are symporters of 1 H^+^ : 1 K^+^ stoichiometry.
7. A steady state condition of 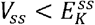 is (indirect) proof of functional H^+^/K^+^ antiporters in the membrane. The condition 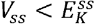 is only achievable with *g*_*HKa*_ > 0.
8. Most cost efficient are antiporters of 1 H^+^ : 1 K^+^ stoichiometry, i.e. they function as electroneutral antiporters.
9. Hyperpolarization-activated K_in_ channels may serve in homeostatic conditions as K^+^ release channels when acting together with H^+^/K^+^ symporters.
10. Depolarization-activated K_out_ channels may serve in homeostatic conditions as K^+^ uptake channels when acting together with H^+^/K^+^ antiporters.

It might be surprising that all these far-reaching conclusions could be drawn solely from the knowledge on the thermodynamics of the different transport processes (Δ*μ*). More knowledge about the transporters (e.g. K_M_-values or voltage-dependence) was ***<u>not</u>*** necessary. Actually, all these details are covered by the *g*_*X*_ values. However, this finding indicates the enormous robustness of the biological system. The basic principle of homeostasis (steady state) does not depend on molecular details but shows a universal pattern that can be achieved in a huge variety of different nuances. It is therefore highly tolerant towards disturbances, such as many mutations in the transporter proteins and could be maintained even in dramatic evolutionary processes. It is thus not individual transporters but rather transporter networks (homeostats) that govern membrane transport and associated homeostatic processes in plant cells.

Conclusions (5-8) are of particular interest. The modeling of the K homeostat points to the existence of both, H^+^/K^+^ symporters and H^+^/K^+^ antiporters. (a) 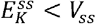 at steady state implies the participation of active H^+^/K^+^ symporters in the homeostat, while (b) 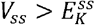 indicates the presence of active H^+^/K^+^ antiporters. A condition of *V* < *E*_*K*_ is not a temporary exception, but is observed quite frequently in plant cells (Thiel et al., 1992; Roelfsema et al., 2001; Roelfsema et al., 2002; Koers et al., 2011). It might be that these cells were not in steady state but in an ongoing process of K^+^ accumulation. However, without H^+^/K^+^ antiporters, the steady state would always be 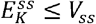, which could lead to rather large K^+^ gradients, especially if the external K^+^ concentration is in the mM range, as is often the case in tissues outside the roots. In such conditions, H^+^/K^+^ antiporters would serve as security valves against an overaccumulation of K^+^. Nevertheless, the role of this type of transporters is largely underrepresented in literature. In comparison to our very good knowledge about K^+^ channels and K^+^/H^+^ symporters, the information about K^+^/H^+^ antiporters in plants is rudimentary (see e.g. Isayenkov et al., 2020 for a recent review). The analyses presented in this study may explain why. Functionally, there was no difference between electroneutral (*n*_*a*_ = 1) and electrogenic (*n*_*a*_ > 1) K^+^/H^+^ antiporters. In the physiological range, both accomplished the same tasks. Nevertheless, systems with electrogenic K^+^/H^+^ antiporters consumed more energy than systems with electroneutral antiporters indicating that optimized organisms like plants would likely employ the electroneutral form. Electroneutral transporters, however, are hardly detectable in standard electrophysiological experiments and therefore extremely difficult to characterize. Thus, the mere fact that *V* < *E*_*K*_ is often observed in a plant cell strongly suggests the existence of electroneutral K^+^/H^+^ antiporters on the basis of the thermodynamic considerations in this study.

It was surprising for us how many fundamental conclusions can be drawn in an airtight manner by applying basic thermodynamics and mathematics to plant biology. The discovery of homeostats enabled a new perspective on membrane transport in plants. And with the considerations presented here for K homeostats, we may have facilitated to adapt this concept to any type of homeostat as already carried out for anions (Dreyer et al., 2022; Li et al., 2024), sugars (Dreyer, 2021a) and auxins (Geisler and Dreyer, 2024), for instance. In a follow-up tutorial study we will present the analyses of the dynamics of the K homeostat. i.e. how the system reacts when it is deflected from the steady state. We will present the mathematical background that describes when the K homeostat is brought out of steady state (i) by a readjustment of transporter activities (tuning of *g*_*X*_ values), (ii) by an external imbalance in the K^+^ gradient, or (iii) by an electrical stimulus. Here, too, the detour via mathematics and computer simulations allows far-reaching conclusions to be drawn about the fundamental physiological properties of the transporter network.

## Author Contributions

ID conceived the study. All authors contributed to the design of the study. ID conducted the data analyses. NH-R, YB-H, VT-C, SZA-M, ED-P, FM-Q, FV-V, OA-S, and MER-M participated in data visualization. ID, JR, and EM wrote the first draft of the manuscript. All authors contributed to the editing of the manuscript and approved its final version.

## Financial Support

This work was supported by the Agencia Nacional de Investigación y Desarrollo de Chile (ANID), grant No. 21220432 to F.M.-Q., No. 21220419 to F.V.-V., and Anillo-ANID ATE220043 (the multidisciplinary center for biotechnology and molecular biology for climate change adaptative in forest resources; CeBioClif) to I.D. and by Fondo Nacional de Desarrollo Científico, Tecnológico y de Innovación Tecnológica (FONDECYT/Chile), grant No. 3190544 to M.E.R.-M., No. 1210920 to E.M, and No. 1220504 to I.D.

## Conflicts of Interest

None.

## Data and Coding Availability Statement

The authors confirm that the data supporting the findings of this study are available within the article and its supplementary materials. Raw data sets are available from the corresponding author, ID, upon reasonable request.

## Notes

### Competing Interest Statement

The authors have declared no competing interest.

## References

Anschütz U, Becker D, Shabala S (2014) Going beyond nutrition: Regulation of potassium homoeostasis as a common denominator of plant adaptive responses to environment. J Plant Physiol 171: 670–687

Blatt MR (2024) A charged existence: A century of transmembrane ion transport in plants. Plant Physiol. doi: 10.1093/PLPHYS/KIAD630

Britto DT, Coskun D, Kronzucker HJ (2021) Potassium physiology from Archean to Holocene: A higher-plant perspective. J Plant Physiol 262: 153432

Carpaneto A, Geiger D, Bamberg E, Sauer N, Fromm J, Hedrich R (2005) Phloem-localized, proton-coupled sucrose carrier ZmSUT1 mediates sucrose efflux under the control of the sucrose gradient and the proton motive force. Journal of Biological Chemistry 280: 21437–21443

Dreyer I (2017) Plant potassium channels are in general dual affinity uptake systems. AIMS Biophys 4: 90–106

Dreyer I (2021a) Nutrient cycling is an important mechanism for homeostasis in plant cells. Plant Physiol 187: 2246–2261

Dreyer I (2021b) Potassium in plants – Still a hot topic. J Plant Physiol 261: 153435

Dreyer I, Blatt MR (2009) What makes a gate? The ins and outs of Kv-like K +channels in plants. Trends Plant Sci 14: 383–390

Dreyer I, Li K, Riedelsberger J, Hedrich R, Konrad KR, Michard E (2022) Transporter networks can serve plant cells as nutrient sensors and mimic transceptor-like behavior. iScience 25: 104078

Dreyer I, Michard E (2020) High- and low-affinity transport in plants from a thermodynamic point of view. Front Plant Sci 10: 1797

Dreyer I, Sussmilch FC, Fukushima K, Riadi G, Becker D, Schultz J, Hedrich R (2021) How to Grow a Tree: Plant Voltage-Dependent Cation Channels in the Spotlight of Evolution. Trends Plant Sci 26: 41–52

Dreyer I, Uozumi N (2011) Potassium channels in plant cells. FEBS Journal 278: 4293–4303

Epstein E, Hagen CE (1952) A kinetic study of the absorption of alkali cations by barley roots. Plant Physiol 27: 457–474

Epstein E, Rains DW, Elzam OE (1963) Resolution of dual mechanisms of potassium absorption by barley roots. Proc Natl Acad Sci U S A 49: 684–692

Geisler M, Dreyer I (2024) An auxin homeostat allows plant cells to establish and control defined transmembrane auxin gradients. bioRxiv 2024.02.07.579341

Hedrich R (2012) Ion channels in plants. Physiol Rev 92: 1777–1811

Hille B (2001) Ion channels of excitable membranes, 3rd ed. Ion channels of excitable membranes. doi: 10.1007/3-540-29623-9_5640

Isayenkov S V., Dabravolski SA, Pan T, Shabala S (2020) Phylogenetic diversity and physiological roles of plant monovalent cation/H+ antiporters. Front Plant Sci 11: 1

Jegla T, Busey G, Assmann SM (2019) Evolution and Structural Characteristics of Plant Voltage-Gated K+ Channels. Plant Cell 30: 2898–2909

Koers S, Guzel-Deger A, Marten I, Roelfsema MRG (2011) Barley mildew and its elicitor chitosan promote closed stomata by stimulating guard-cell S-type anion channels. The Plant Journal 68: 670–680

Li K, Grauschopf C, Hedrich R, Dreyer I, Konrad KR (2024) K+ and pH homeostasis in plant cells is controlled by a synchronized K+/H+ antiport at the plasma and vacuolar membrane. New Phytologist. doi: 10.1111/NPH.19436

Lohse G, Hedrich R (1992) Characterization of the plasma-membrane H+-ATPase from Vicia faba guard cells - Modulation by extracellular factors and seasonal changes. Planta 188: 206–214

Nour-Eldin HH, Andersen TG, Burow M, Madsen SR, Jørgensen ME, Olsen CE, Dreyer I, Hedrich R, Geiger D, Halkier BA (2012) NRT/PTR transporters are essential for translocation of glucosinolate defence compounds to seeds. Nature 488: 531–534

Reyer A, Häßler M, Scherzer S, Huang S, Pedersen JT, Al-Rascheid KAS, Bamberg E, Palmgren M, Dreyer I, Nagel G, et al (2020) Channelrhodopsin-mediated optogenetics highlights a central role of depolarization-dependent plant proton pumps. Proceedings of the National Academy of Sciences 117: 20920–20925

Rienmüller F, Dreyer I, Schönknecht G, Schulz A, Schumacher K, Nagy R, Martinoia E, Marten I, Hedrich R (2012) Luminal and cytosolic pH feedback on proton pump activity and ATP affinity of V-type ATPase from Arabidopsis. Journal of Biological Chemistry 287: 8986–8993

Roelfsema MRG, Hanstein S, Felle HH, Hedrich R (2002) CO2 provides an intermediate link in the red light response of guard cells. The Plant Journal 32: 65–75

Roelfsema MRG, Steinmeyer R, Staal M, Hedrich R (2001) Single guard cell recordings in intact plants: Light-induced hyperpolarization of the plasma membrane. Plant Journal 26: 1–13

Sharma T, Dreyer I, Riedelsberger J (2013) The role of K channels in uptake and redistribution of potassium in the model plant Arabidopsis thaliana. Front Plant Sci 4: 224

Stanton C, Sanders D, Krämer U, Podar D (2022) Zinc in plants: Integrating homeostasis and biofortification. Mol Plant 15: 65–85

Thiel G, MacRobbie EAC, Blatt MR (1992) Membrane transport in stomatal guard cells: The importance of voltage control. J Membr Biol 126: 1–18

Ward JM, Mäser P, Schroeder JI (2009) Plant Ion Channels: Gene Families, Physiology, and Functional Genomics Analyses. Annu Rev Physiol 71: 59–82

