## Supplementary material for "Homeostats – the hidden rulers of ion homeostasis in plants": Figure S1

### Supplementary Figure S1

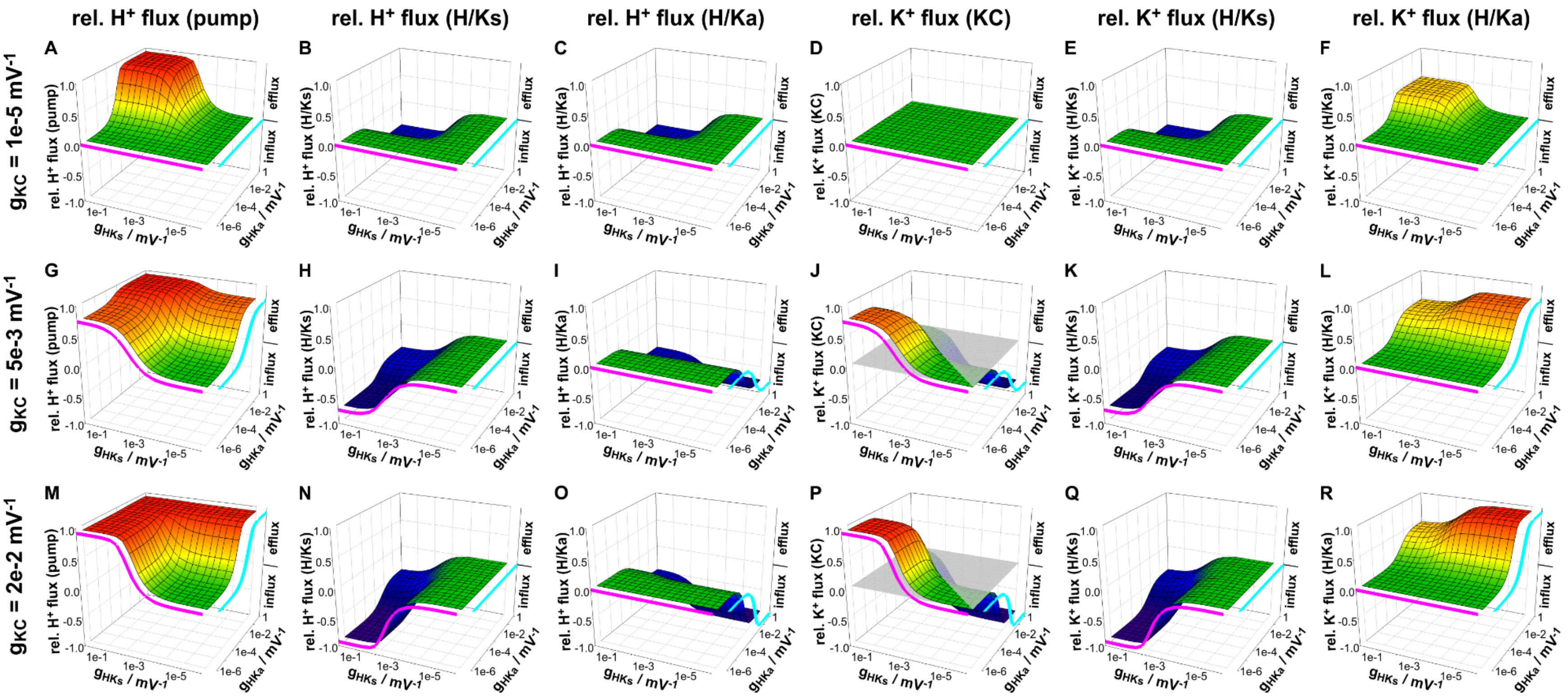


**Figure S1. $H^+$ and $K^+$ fluxes through the different transporters in homeostatic (steady state) conditions.** Dependency of the $H^+$ and $K^+$ fluxes on the activities of the $K^+$ channels ($g_{KC}$), $H^+/K^+$ symporters ($g_{HKs}$) and $H^+/K^+$ antiporters ($g_{HKa}$). (**A, G, M**) Relative $H^+$ flux mediated by the $H^+$ ATPase. (**B, H, N**) Relative $H^+$ flux mediated by the $H^+/K^+$ symporter. (**C, I, O**) Relative $H^+$ flux mediated by the $H^+/K^+$ antiporter. (**D, J, P**) Relative $K^+$ flux mediated by the $K^+$ channel. (**E, K, Q**) Relative $K^+$ flux mediated by the $H^+/K^+$ symporter. (**F, L, R**) Relative $K^+$ flux mediated by the $H^+/K^+$ antiporter. The fluxes are shown relative to the maximal $H^+$ efflux that can be generated by the $H^+$ ATPase ($J_{Hmax} = I_{Hmax}/e_0$). Data were calculated for the case $n_s = 1$, $n_a = 1$, $V_{0,pump}$ = -200 mV, and $E_H$ = +57.6 mV ($\Delta$pH = 1). The magenta lines show the values in the absence of active $H^+/K^+$ antiporters ($g_{HKa}$ = 0), whereas the cyan lines indicate the values in the absence of active $H^+/K^+$ symporters ($g_{HKs}$ = 0).
